# A novel bioinformatics approach reveals key transcription factors regulating type 1 diabetes-associated transcriptomes

**DOI:** 10.1101/2025.05.13.653885

**Authors:** Afzal Sheikh, Sultana Parvin, Shalman Dipto, Shamim Hossain

## Abstract

We developed a novel bioinformatics pipeline that reveals key transcription factors (TFs) regulating type 1 diabetes transcriptomes by combining automated Python workflows, DESeq2-based differential expression analysis, motif enrichment, and genome-wide TF abundance profiling, a rare but powerful strategy that deepens our understanding of disease mechanisms. This approach not only uncovers TFs driving pathology but also quantifies TF abundance across multiple genomic loci, enabling rapid and precise monitoring of their occupancy. Analyzing 21 RNA sequencing datasets, including 13 from early-stage T1D patients and 8 matched controls, we identified nearly 6 000 differentially expressed genes; 1 900 met strict significance and fold-change criteria and 211 are previously uncharacterized transcripts that distinguish T1D from healthy samples. Pathway analysis highlighted disruptions in beta-cell signaling, ALK-linked drug responses, neurodegenerative processes, and cytoskeletal organization. Upstream motif analysis revealed enrichment of Myc/Max, AP-1, SP-1, TATA-box, and NF-κB binding sites in upregulated genes, confirming their central role in the T1D transcriptome. By placing TFs at the core of our discovery platform, this work uncovers novel molecular drivers of T1D and identifies actionable biomarkers and therapeutic targets for personalized treatment.

## Introduction

Type 1 diabetes (T1D) is an autoimmune disorder characterized by the destruction of insulin-producing beta cells in the pancreas, leading to insulin deficiency and chronic hyperglycemia (1). Unlike Type 2 Diabetes (T2D), which is primarily driven by insulin resistance, T1D is caused by an immune-mediated attack, often triggered by environmental factors in genetically predisposed individuals (2,3). Although the exact cause of this disease remains unclear, both genetic and environmental factors contribute to its onset. The onset of the disease typically occurs in childhood or adolescence, although adult-onset cases are increasingly recognized (4,5). This chronic condition requires lifelong insulin therapy and close monitoring of blood glucose levels to prevent complications, including diabetic ketoacidosis (DKA), organ failure, cardiovascular disease, neuropathy, retinopathy, and kidney disease and death (6–9).

T1D management primarily relies on exogenous insulin therapy, administered through multiple daily injections (MDI) or continuous subcutaneous insulin infusion (CSII) via insulin pumps, aiming to replicate physiological insulin secretion (10). Advancements in insulin analogs—rapid-acting, long-acting, and ultra-long-acting have enhanced glycemic control and minimized hypoglycemia risk (11,12). Additionally, technological innovations like continuous glucose monitoring (CGM) and hybrid closed-loop insulin delivery systems (artificial pancreas) have further optimized real-time glucose management (13). Beyond insulin, adjunctive therapies like sodium-glucose cotransporter-2 (SGLT2) inhibitors and glucagon-like peptide-1 (GLP-1) agonists are being studied to enhance metabolic control in T1D (14). Immunomodulators (e.g., teplizumab), islet transplantation, and stem cell therapies also show promise for preserving β-cell function or achieving remission (15, 16). Despite advancements, T1D remains a lifelong challenge due to persistent risks of hypoglycemia/hyperglycemia, variable insulin absorption, and the burden of continuous management (17). Some patients develop insulin resistance (“double diabetes”), particularly those with obesity, metabolic syndrome, or prolonged disease duration, highlighting the need for research to address variable treatment efficacy and safety concerns (18–20).

Although clinical management of T1D has evolved with improved glycemic monitoring and insulin therapies, the underlying transcriptomic changes that precede and accompany disease onset remain poorly understood. Increasing evidence suggests that early molecular alterations in gene expression and regulatory elements may play critical roles in autoimmunity, beta-cell dysfunction, and immune system dysregulation in T1D patients (21–25). However, most studies have focused on bulk phenotypic or immunological features, with limited resolution on gene-level regulatory mechanisms. Notably, few have explored novel gene transcripts, and transcription factor binding patterns, promoter-linked regulatory hotspots in human peripheral blood cells during early T1D progression (25, 26). These knowledge gaps hinder the identification of early biomarkers and actionable targets for precision therapies. Thus, transcriptomic profiling of human white blood cells (WBCs) provides a promising strategy to uncover T1D-specific molecular signatures and novel regulatory pathways. These insights are especially critical, given the variable efficacy and potential safety concerns associated with current treatment approaches, underscoring the need for ongoing research to develop more effective and sustainable T1D management strategies.

With the advent of high-throughput technologies, transcriptomics which uses RNA sequencing data has emerged as a powerful tool for investigating the genetic factors, environmental life-style related cause including its molecular mechanisms underlying T1D pathogenesis (27, 28).

A critical component of this approach is the application of pathway enrichment analyses, such as those provided by the Kyoto Encyclopedia of Genes and Genomes (KEGG) and Gene Ontology (GO), which facilitate the identification of biological processes, molecular functions and cellular components aberrantly regulated in disease states (29). The implications of these pathway alterations extend beyond our understanding of disease mechanisms; they also highlight potential targets for therapeutic intervention. The convergence of transcriptomic data with pathway enrichment analyses underscores the intricate interplay between gene regulation, metabolic control, and cellular signaling. By dissecting these complex networks, researchers can uncover novel targets including the novel genes with unknow functions for intervention and develop strategies that are more precisely tailored to correct the multifaceted disruptions observed in metabolic disorders particularly T1D patients. As the field continues to evolve, the integration of these molecular insights with clinical research holds promise for the development of next-generation therapeutics that can more effectively address the underlying causes of disease (1, 30–31).

Although RNA-Seq and whole-genome analyses have catalogued extensive disease-associated gene repertoires, the functional contribution of transcription factors (TFs) to disease-specific transcriptomic architectures remains under-investigated. TFs act as master regulators of gene expression, and quantitative profiling of their genomic occupancy is indispensable for elucidating pathogenic mechanisms and informing therapeutic design. To address this gap, we have developed an integrative bioinformatics platform that couples automated Python-based preprocessing, DESeq2-mediated differential expression analysis, de novo motif enrichment, and quantitative assessment of TF abundance across multiple genomic loci. Application of this pipeline to secondary RNA-Seq data from 13 newly diagnosed T1D patients and eight age- and sex-matched healthy controls enabled the simultaneous identification of novel, disease-segregating transcripts and precise quantification of key TFs. Our findings delineate the TFs that orchestrate T1D-specific transcriptomic reprogramming and establish a foundation for TF-centric therapeutic strategies aimed at restoring normative gene regulatory networks in Type 1 diabetes.

## Materials and Methods

### Secondary Data selection

We analyzed secondary RNA-Seq data from whole blood samples collected within 48 hours of diagnosis from 13 new-onset Type 1 Diabetes (T1D) patients and 8 age-, sex-, and BMI-matched healthy controls of diagnosis at Riley Hospital for Children, with parental consent and participant assent (ages >7 years). A total of 21 RNA-Seq datasets were randomly selected from the National Library of Medicine’s Sequence Read Archive (SRA) from the study RNA-seq data published by Webb-Robertson et al. (32). The raw SRA files were downloaded and converted into FASTQ format using standard bioinformatics tools. The library construction protocol, including total RNA extraction cDNA synthesis, library quality checks etc., downstream analysis flow has been described in the above study. RNA sequencing data are publicly available in the NCBI Sequence Read Archive (SRA) under BioProject accession SRP532541, (accession link: https://www.ncbi.nlm.nih.gov/Traces/study/?acc=SRP532541&o=acc_s%3Aa).

Raw sequencing files for control (4 from cohort 1 and 4 from cohort 2) include: SRR30660548, SRR30660551, SRR30660552, SRR30660559, SRR30660588, SRR30660589, SRR30660590, SRR30660591, and T1D (7 from cohort 1 and 6 from cohort 2): SRR30660553-55, SRR30660560-62, SRR30660572-74, SRR30660577, SRR30660582, SRR30660583, SRR30660584.

### Data processing

In our RNA-seq analysis pipeline, raw sequencing data (SRR30660XXX) were downloaded from GEO using the NCBI SRA Toolkit’s prefetch utility (v.3.0, default settings to ensure reliable retrieval). The downloaded SRA files were then converted to paired-end FASTQ format with fastq-dump --split-files --gzip, splitting reads into forward and reverse files and compressing them to enhance storage efficiency by SRAtoolkit. Adapter and quality trimming were performed using Trimmomatic v0.39 with parameters TRAILING:10, SLIDINGWINDOW:4:15, and MINLEN:36 to remove low-quality bases and short fragments, thus improving downstream alignment accuracy. Trimmed reads were aligned to the human reference genome (GRCh38) using HISAT2 v2.2.1 with --dta for downstream transcriptome assembly, -p 4 threads, and default mismatch penalties (MX=6, MN=2) optimized for spliced alignment as per the HISAT2/Tool Shed manual. The resulting SAM files were converted to BAM via samtools view -bS and coordinate-sorted with samtools sort -o, producing sorted BAM files ready for quantification as per the samtools manual. Gene-level counts were obtained using featureCounts (Subread v2.0.3) with -p (paired-end), -s 2 (strand-specific), -T 4 threads, and an Ensembl GTF annotation to assign reads accurately to gene features (ShiLab-Bioinformatics/subread-GitHub). These count matrices were merged in Python, and gene lengths were used to compute RPKM and TPM according to the formulas RPKM = (read count × 10^9)/(gene length × total reads) and TPM = (RPKM/sum RPKM) × 10^6 for proper within- and between-sample normalization. For differential expression analysis, DESeq2’s median-of-ratios normalization was applied to correct for library size and composition bias, followed by dispersion estimation and Wald testing under a negative-binomial model, differential expression analysis using DESeq2. Finally, p-values were adjusted using the Benjamini–Hochberg procedure to control the false discovery rate at p = 0.05, ensuring robust identification of differentially expressed genes. An automated python script was generated to run all the above tools with specified functions as per their manual.

### Gene Expression Analysis using R

Differential expression analysis was performed using DESeq2 (v1.42.1 in R (v4.5.0) on RNA-seq raw counts from control and type 1 diabetes (T1D). Raw count datasets were merged, missing values replaced with zero, and duplicate gene counts aggregated to ensure reproducibility. A DESeq2 dataset object was constructed with integrated sample metadata, followed by normalization and dispersion estimation. Log-transformed expression values (rlog) were then visualized using various applications packages, facilitating downstream interpretation. Differential expression testing between T1D and control groups was conducted using default negative binomial generalized linear models, with significance thresholds set at an adjusted p-value < 0.05. The R scripts used for data analysis and visualization are available upon reasonable request.

### Minus vs. Average plot (‘M’ represents the log ratio and ‘A’ represents Average)

MA plot was generated to visualize log₂ fold changes (M) against mean expression intensity (A). Genes were classified as follows: (1) log₂FC > 2 (upregulated) or < –2 (downregulated), highlighted in red and blue, respectively; (2) genes with highly differential expression (log₂FC > 5 or < –5), representing the highest magnitude of transcriptional changes, were distinctly marked with dark red (upregulated) or dark blue (downregulated) symbols. Statistically significant genes (p < 0.05) were further annotated with orange (upregulated) or yellow (downregulated) borders to distinguish them from non-significant, yet highly variable, candidates. These visualizations were optimized to emphasize both the magnitude and statistical confidence of transcriptional alterations associated with T1D pathogenesis.

### Principal component analysis (PCA)

Principal component analysis was conducted on the transformed count data using rlog() for variance stabilizing transformation. The results were visualized in a PCA plot, with the percentage of variance explained by the first two principal components indicated on the axes.

### Heatmap of Top Differentially Expressed Genes

A heatmap of all the significantly and differentially expressed genes (ranked by adjusted p-value) was generated using the pheatmap () function. The heatmap showed clustered rows and columns based on gene expression profiles.

### KEGG and GO Pathway Enrichment Analysis

Post-normalization, DESeq2 results were mapped from ENSEMBL to Entrez IDs using the org.Hs.eg.db package. KEGG pathway enrichment was performed using the clusterProfiler package’s enrichKEGG() function (p < 0.05), and GO enrichment analysis was performed using enrichGO() (ontology: Biological Process, p-value cutoff = 0.05). Enrichment results were visualized via dot plots generated by the enrichplot and ggplot2 packages.

### Reactome Pathway Analysis

An overrepresentation analysis was conducted using Reactome pathway analysis with a hypergeometric distribution test to determine enrichment of specific pathways. All p-values were adjusted for multiple comparisons using the Benjamini-Hochberg method. The analysis was restricted to human data by converting non-human identifiers to their human equivalents, ensuring that only Homo sapiens pathways were considered.

### High-Throughput Analysis of Transcription Factor Binding Sites in Human Promoters

Using a human-robust pipeline, first-exon plus 5 kb upstream and 2 kb downstream sequences for each Ensembl gene ID were retrieved via the Ensembl REST API in Python with built-in retry logic, filtering for protein-coding biotypes and excluding non-canonical contigs to avoid HTTP errors (33–34). IUPAC-encoded TF motifs were converted into regular-expression patterns and scanned for both exact and ∼10% fuzzy matches using the regex module’s approximate-matching flags PyPI. All binding-site hits were compiled into CSV files and ingested via pandas.read_csv for efficient tabular manipulation Pandas (35). Per-gene multi-panel figures—comprising scatter plots, stacked density histograms via seaborn.histplot, bar charts, lollipop diagrams, and ECDFs— were generated with Matplotlib and Seaborn. Individual TIFF/JPEG figures were bundled into multi-page PDFs using matplotlib.backends.backend_pdf.PdfPages for publication-ready reports Matplotlib. Finally, comparative summary barplots and heatmaps contrasting up-versus down-regulated gene sets were rendered via Seaborn’s barplot and heatmap, with upstream download, scanning, and plotting stages parallelized using concurrent.futures ThreadPoolExecutor and ProcessPoolExecutor for I/O and CPU efficiency.

## Results

### Differential expression profiling identifies 5,840 DEGs and distinct transcriptomic signatures in type 1 diabetes

In our analysis, a total of 5,840 differentially expressed genes (DEGs) were identified in T1D patients compared to controls, including 4,800 upregulated and 1,040 downregulated transcripts. Applying stringent criteria (p < 0.05, |log₂FC| > 2), 1,916 robust DEGs were retained. Among upregulated genes, 368 met significance thresholds, while 533 exhibited higly upregulation (log₂FC > 5; 221 significant). Downregulated genes included 96 significant candidates, with 44 showing highly downregulation (log₂FC < –5; 19 significant). Moreover, our analysis identified 164 significant novel transcripts within the upregulated set and 47 among the downregulated genes (Table_1), highlighting potential disease-associated targets.

The MA plot (Figure 1A) revealed distinct transcriptional alterations in T1D, with a predominance of upregulated genes indicating widespread gene activation. Principal Component Analysis (PCA) of rlog-transformed data revealed clear separation between T1D and controls, with PC1 (54% variance) and PC2 (12%) capturing 66% of total variance (Figure 1B). This clustering underscores a distinct T1D transcriptional profile, supporting widespread dysregulation linked to pathogenesis.

**Figure 1.**
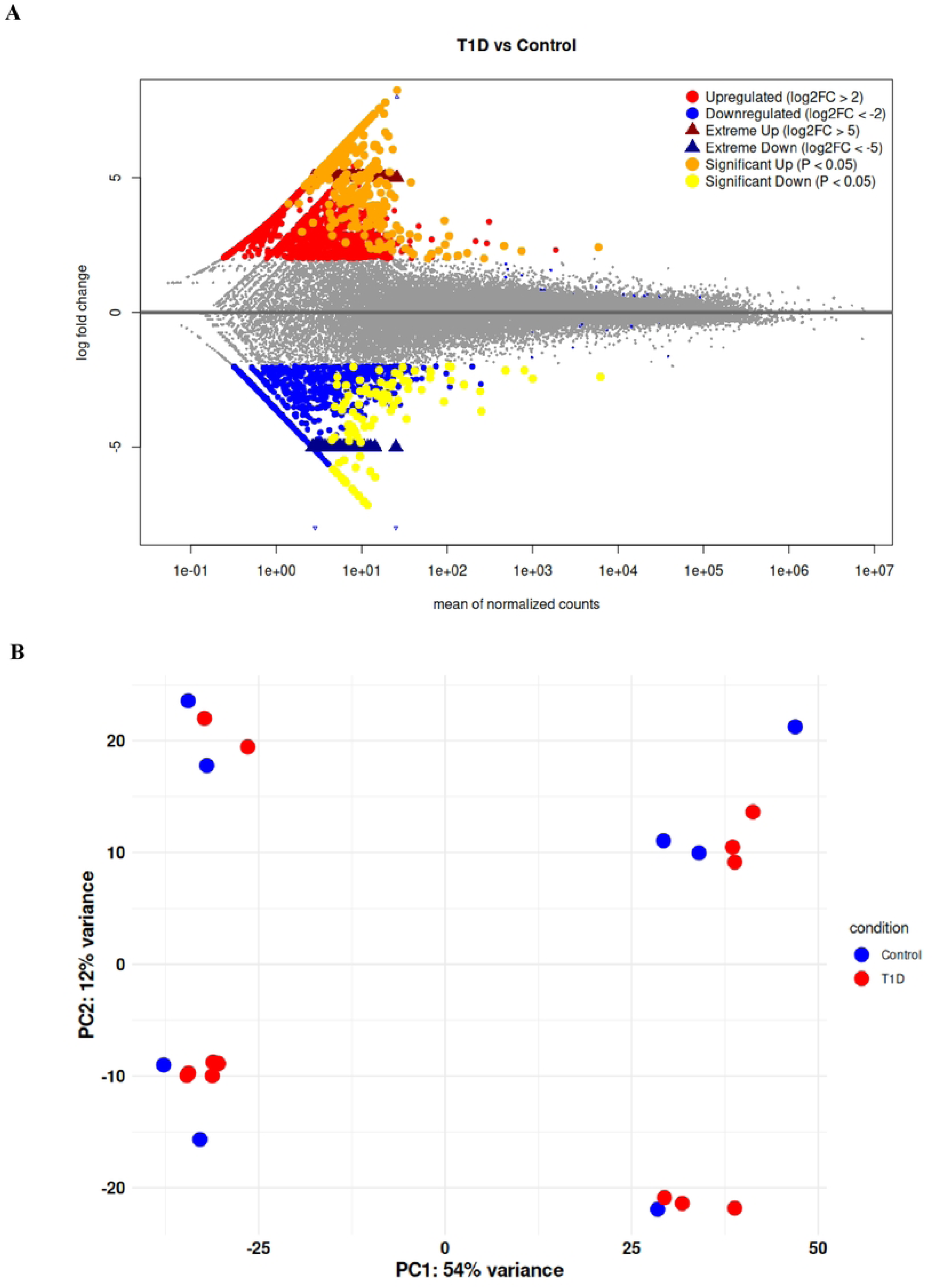
MA plot and PCA reveal differential gene expression and sample clustering in T1D. **(A)** The MA plot depicts differential gene expression between T1D and control samples, with the y-axis represents log₂ fold change while the x-axis shows mean normalized counts. Genes clustering around zero show minimal expression differences, whereas red points denote moderate upregulation (log₂FC > 2), and blue points indicate moderate downregulation (log₂FC < –2). Highly changes (log₂FC > 5 and < –5) are marked by dark red and dark blue triangles, respectively, with statistically significant genes further highlighted in orange (upregulated) and yellow (downregulated). **(B)** Principal Component Analysis (PCA) was performed on rlog-transformed RNA-seq data, with PC1 and PC2 explaining 54% and 12% of the variance, respectively. The clear segregation of T1D and control samples indicates distinct transcriptional profiles, where each point represents an individual sample.

### Integrated transcriptomic profiling and enrichment analyses (GO and KEGG) highlight distinct molecular signatures and dysregulated pathways in T1D

Among the differentially expressed genes, between T1D and controls, 1,916 genes met stringent criteria (p < 0.05 and |log₂FC| > 2) and were visualized using a heatmap (Figure 2A), which revealed distinct clusters: one enriched in genes linked to immune activation, inflammation, and metabolic dysregulation, and another associated with β-cell function, insulin signaling, and cellular homeostasis. Comprehensive KEGG pathway analysis showed significant enrichment in multiple biological pathways, notably neurodegenerative disease pathways with approximately 400 genes in the general category and over 300 genes in pathways for Alzheimer’s disease, ALS, and Huntington’s disease alongside pathways related to focal adhesion, transport and catabolism, endocytosis, cell growth and death, and signal transduction (Figure 2B).

**Figure 2.**
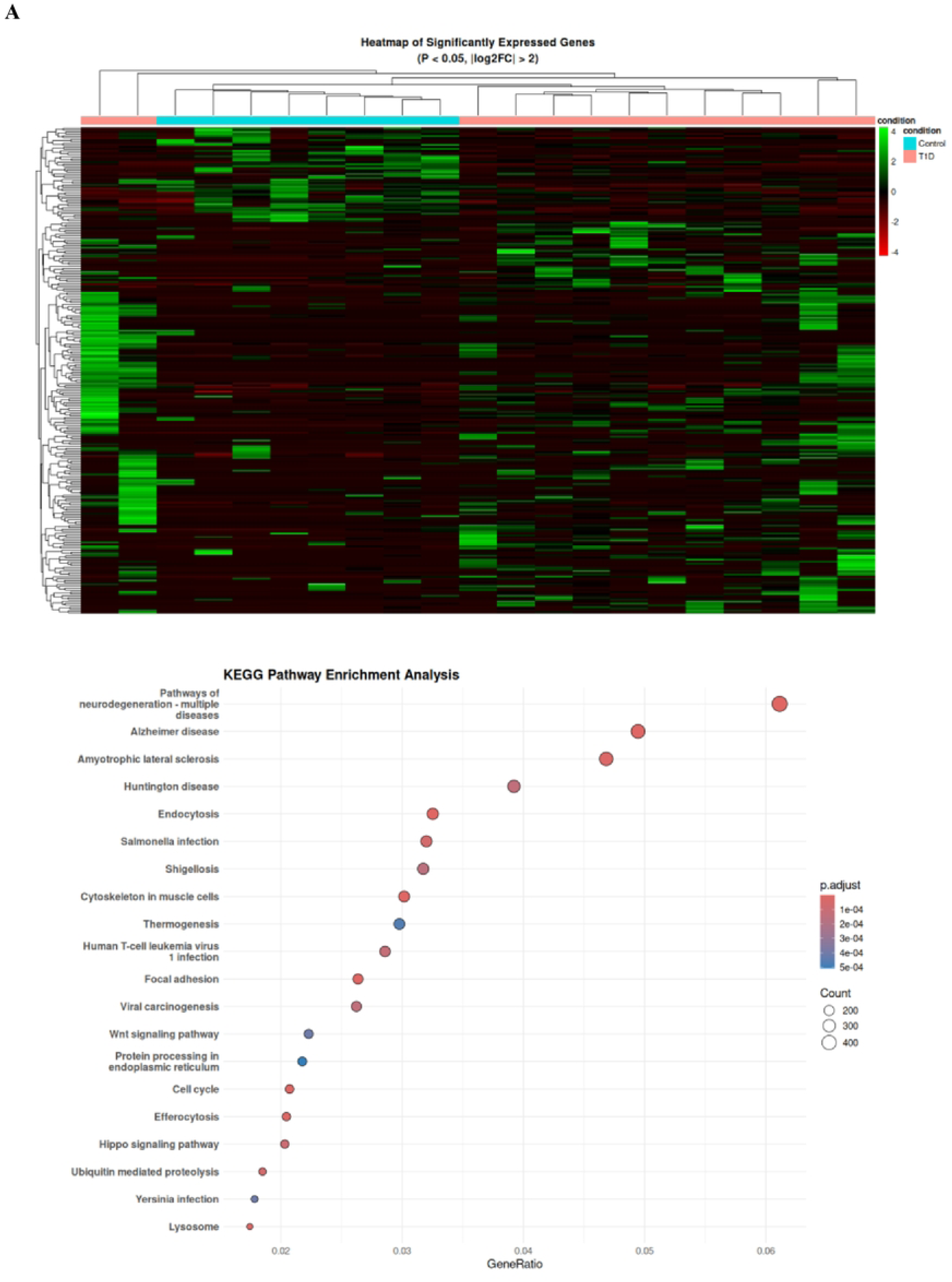
Heatmap of DEGs and KEGG pathway enrichment in T1D. **(A)** Heatmap: Hierarchical clustering of rlog-transformed expression values for 1,978 significantly dysregulated genes (p < 0.05, |log₂FC| > 2) between T1D and control samples. The color gradient—red for upregulation, black for intermediate expression, and green for downregulation— highlights distinct clusters; one enriched in immune activation and metabolic dysregulation and another in β-cell function and insulin signaling. **(B)** KEGG Enrichment Analysis: This panel shows the significant enrichment of multiple pathways in T1D, notably neurodegenerative disease pathways (with gene counts ∼400 for the general pathway and >300 for Alzheimer’s, ALS, and Huntington’s diseases) along with pathways related to beta-cell regulation, cytoskeletal organization, and signal transduction. Additionally, analysis of highlyly upregulated genes (log₂FC ≥ 5, p < 0.05) identified the “Cytoskeleton in Muscle Cells” pathway, exhibiting over six-fold upregulation.

On the other hand, GO enrichment analysis further revealed widespread transcriptomic disruptions across Biological Processes (BP), Molecular Functions (MF), and Cellular Components (CC). In BP (Figure 3A), significant enrichment was observed in immune-related processes such as lymphocyte differentiation, macrophage activation, and DNA repair mechanisms including double-strand break repair and chromosome segregation, as well as in developmental processes like cilium organization and embryonic development. In MF (Figure 3B), key alterations included dysregulation of GTPase regulator activity, nucleoside-triphosphatase regulator activity, and protein serine/threonine kinase activity, alongside notable changes in binding functions such as cadherin and histone binding. Finally, CC analysis (Figure 3C) highlighted significant involvement of structural cellular components, with enrichment in cell-substrate junctions, focal adhesions, microtubules, and nuclear elements such as nuclear specks and the nuclear envelope. These findings underscore a multi-layered transcriptomic reprogramming in T1D, implicating disruptions in immune surveillance, genomic stability, signaling pathways, and cellular architecture.

**Figure 3.**
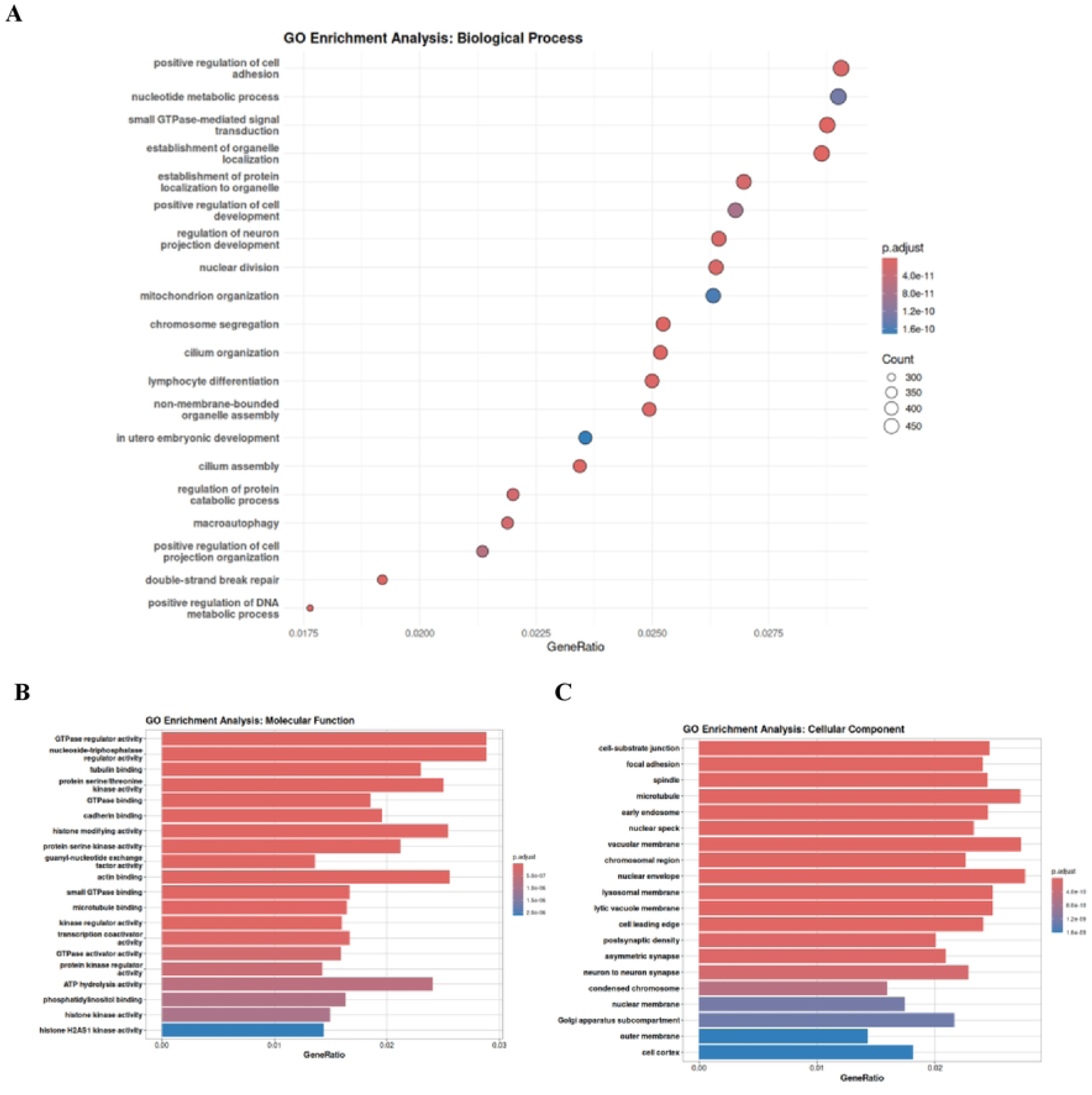
GO enrichment reveals that altered gene expression in T1D affects critical biological processes, molecular functions, and cellular components linked to disease pathogenesis. GO enrichment analysis of significantly differentially expressed genes (log₂FC ≥ 2, p < 0.05) revealed strong enrichment in key functional categories. (A) Dot plot shows the top 20 enriched Biological Processes, including immune response, cell adhesion, metabolic regulation, and signal transduction, indicating transcriptional changes affecting cellular communication and homeostasis in T1D. (B) Molecular Functions such as protein binding, catalytic activity, and ion binding were enriched, reflecting roles in cellular integrity and stress response. (C) Enriched Cellular Components included cell-substrate junctions, focal adhesions, microtubules, and nuclear structures, suggesting widespread gene expression alterations contributing to T1D pathogenesis.

A further detailed examination of highly expression changes revealed that among the 1,916 significant DEGs, 221 genes were highly upregulated (log₂FC ≥ 5, p < 0.05) with 75 of these being highly significantly (p < 0.01) expressed, and 44 genes were highly downregulated (log₂FC < –5), with 19 reaching significance (Table_1., Figure 4).

**Figure 4.**
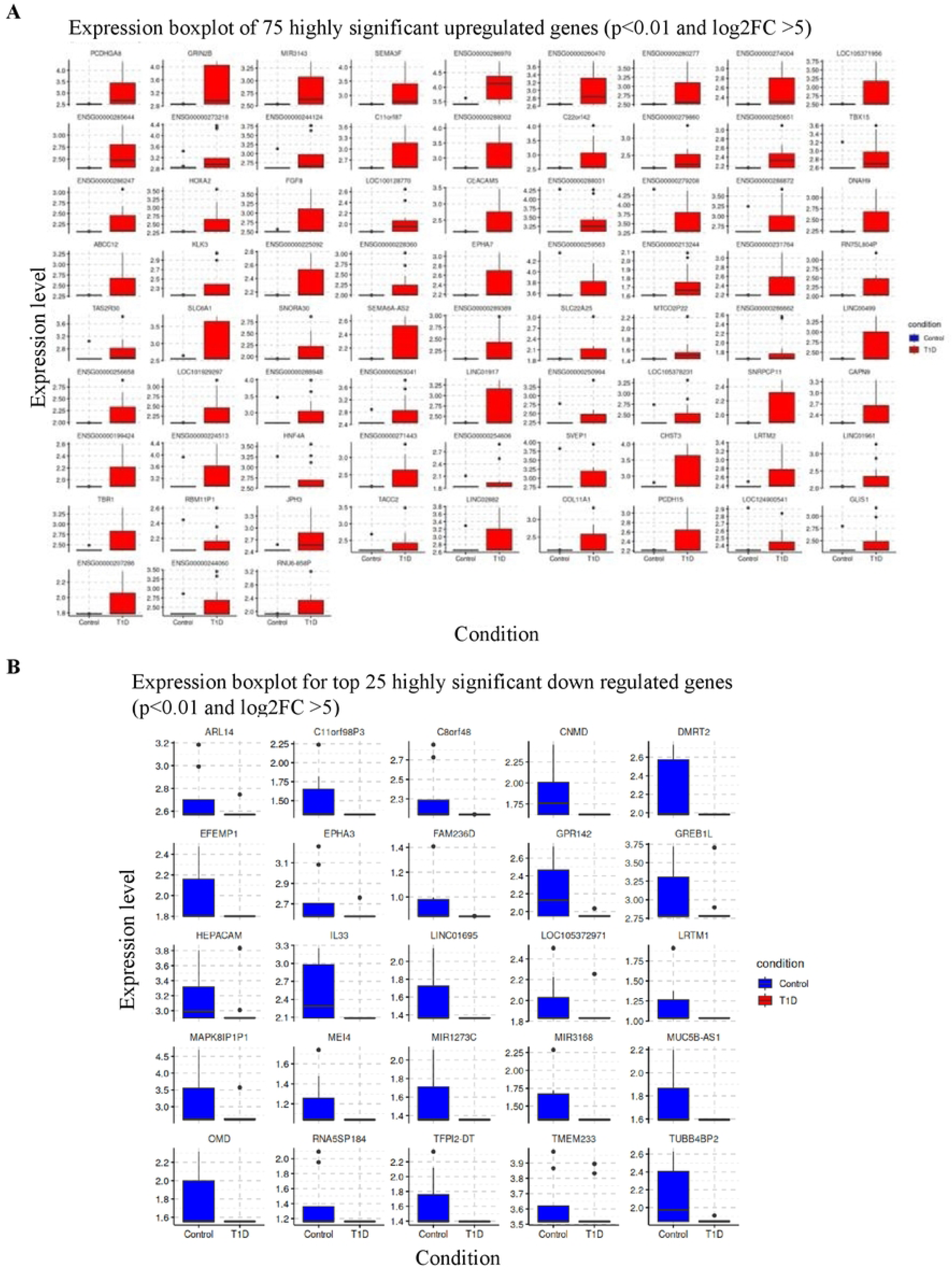
Differential expression of highly significant top 100 up and down regulated genes. **(A)** Highly Upregulated Genes: Displaying 75 genes with log₂ fold changes ≥ 5 and p < 0.01, this panel highlights robust transcriptional activation in T1D compared to controls. **(B)** Highly Downregulated Genes: Featuring 25 genes with log₂ fold changes ≤ -5 and p < 0.01, this panel illustrates pronounced suppression of gene expression in T1D.

### Gene expression patterns in T1D highlights impaired pancreatic β-bell activity and altered immune signaling pathway genes

The transcriptomic analysis revealed that several critical gene sets are significantly down-regulated in T1D patients relative to controls. Notably, genes associated with pancreatic β-cell functions-including AKT2, GCK, PPARGC1A, ATF4, COX4I1, PDX1, UQCRC2, ATPSF1A, IRS1, INSR, MAFA, NDUFS3 and GPX1-were markedly suppressed (Figure 5A and 5B), suggesting impaired insulin secretion and β-cell viability, which are central to the development and progression of T1D (36–38). In parallel, the analysis identified substantial down-regulation of key immune-response genes including CD4, FOXP3, JAK2, NFKB1, STAT1, IFIT3, IL1B, IL6, JAK1, CTLA4, STAT3, IFIT1, TNF, and CCL5 suggesting a marked dysregulation in immune surveillance and inflammatory control that may exacerbate β-cell destruction (Figure 6C and 6D). This finding is in line with previous reports indicating that impaired regulatory T cell function and disrupted cytokine signaling contribute to the pathogenesis of Type 1 Diabetes (39–41). Furthermore, alterations in these signaling pathways have been associated with heightened inflammatory responses and immune dysfunction, which can further accelerate β-cell loss in autoimmune diabetes (23).

**Figure 5.**
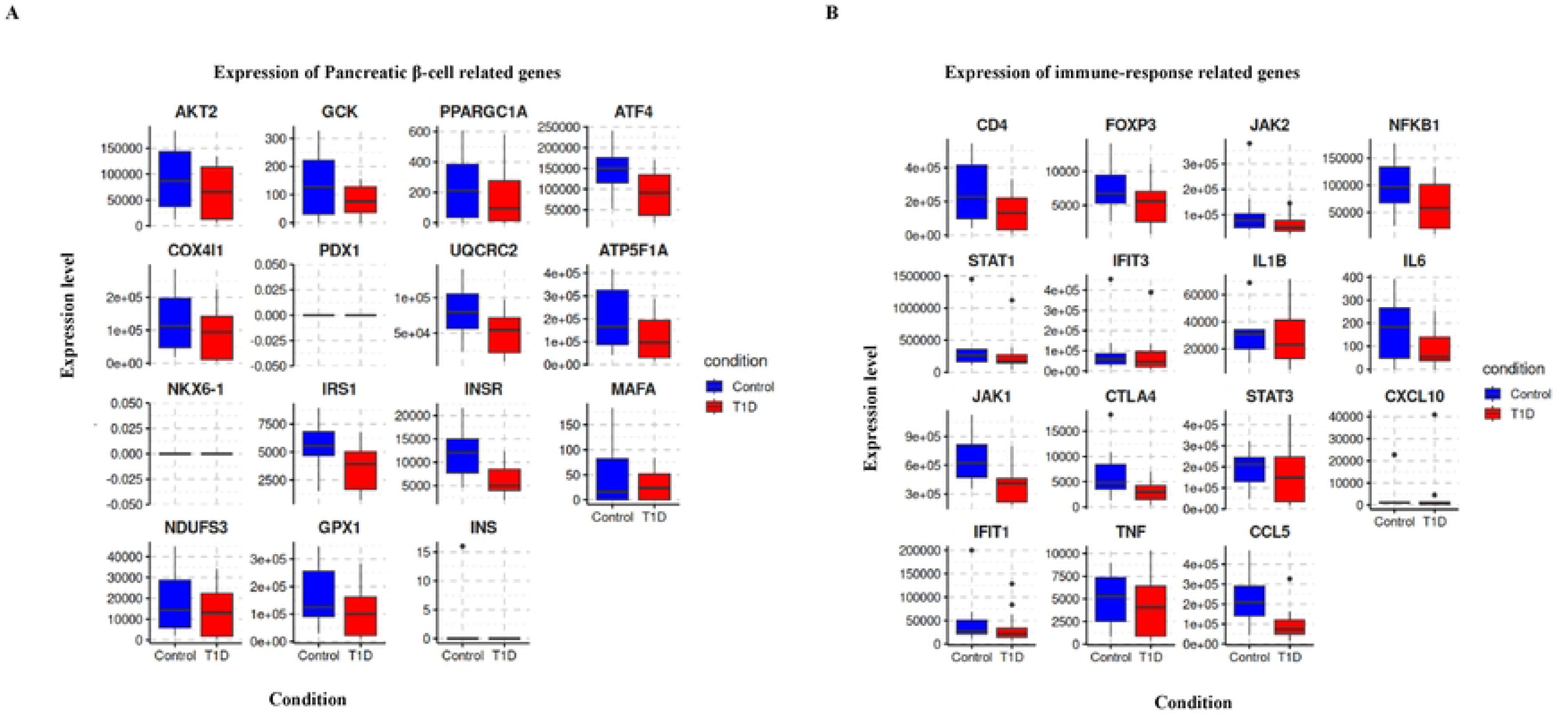
Downregulation of Beta-cell function and immune response genes in T1D. **(A)** Box-Expression Analysis: This panel represents the expression profiles and disease associations of key genes related to pancreatic beta cell function and immune response. Both gene sets are significantly downregulated in T1D compared to controls, suggesting impaired beta cell activity and altered immune regulation.

**Figure 6.**
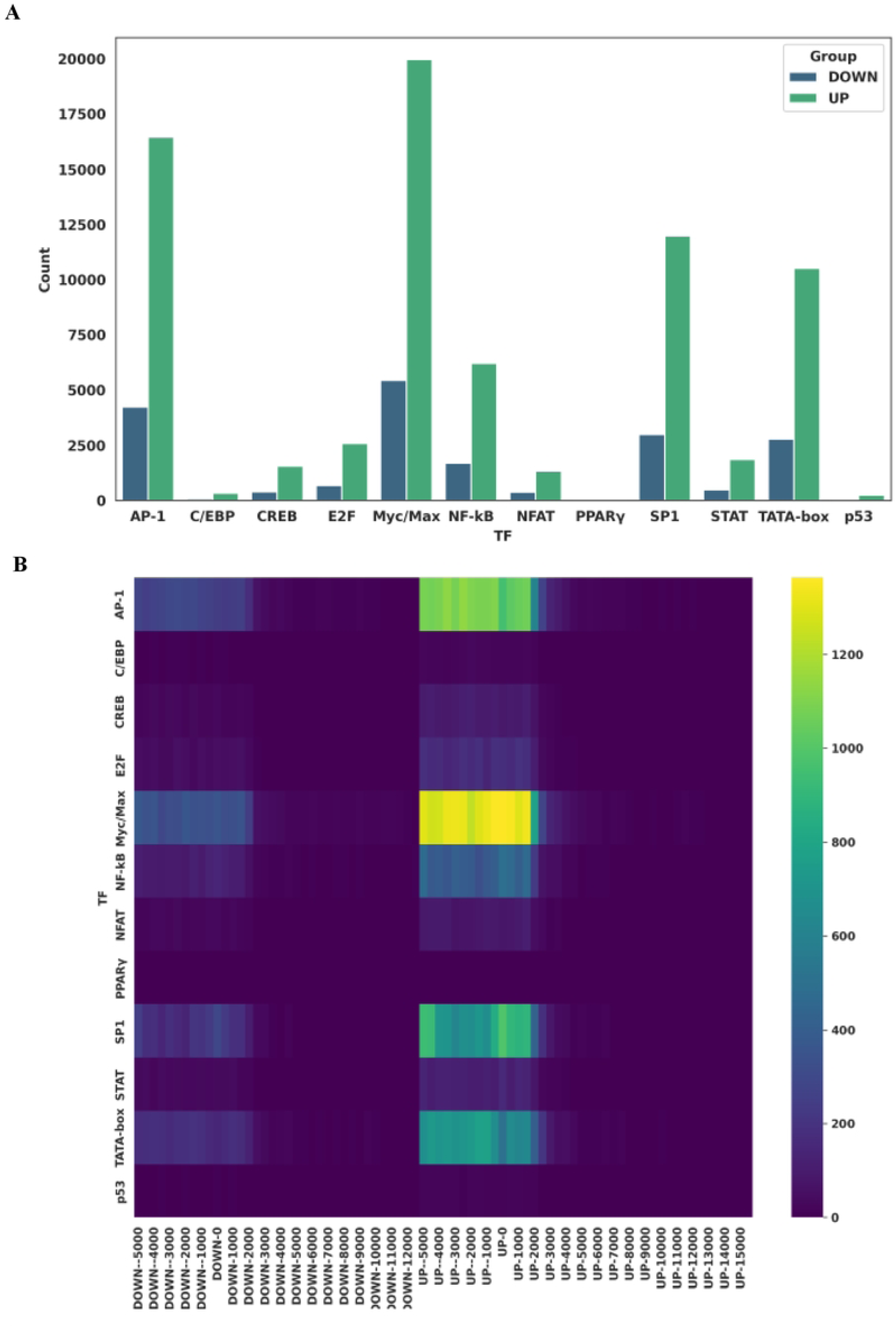
Transcription factor motif enrichment near TSS in T1D DEGs. **(A)** Bar plot showing the number of exact matches for transcription factor (TF) binding motifs within ±5 kb of the transcription start sites of significantly up and down-regulated genes in T1D. A pronounced increase in TF motif occurrences was observed in up-regulated genes, with Myc/Max motifs enriched nearly fivefold, followed by AP-1, SP-1, TATA-box, and NF-κB (∼fourfold). **(B)** Heatmap depicting motif density distribution across the ±5 kb region flanking TSSs, illustrating TF-specific binding site hotspots. Up-regulated transcripts exhibited distinct TF binding clusters compared to down-regulated genes.

### Differentially expressed genes analysis exhibited highly significant novel transcriptomes in T1D patients

Of the 1,916 significantly differentially expressed genes (p < 0.05) in T1D, a subset of 221 highly upregulated genes (log₂FC ≥ 5) was further analyzed, revealing 93 novel transcripts with no gene symbol and known functions. Among these, 44 transcripts lacked any known gene symbols and exhibited highly significant expression (p < 0.001) (Figure 7, Supplement Information_1). Additionally, 183 upregulated genes with known symbols did not map to any Reactome pathway (Supplement Information_2), suggesting the involvement of unidentified pathways or functions in T1D. These findings indicate that, beyond established gene regulatory networks, a reservoir of novel transcripts and unknown gene functions may drive T1D pathogenesis, offering promising avenues for future biomarker discovery and therapeutic intervention.

**Figure 7.**
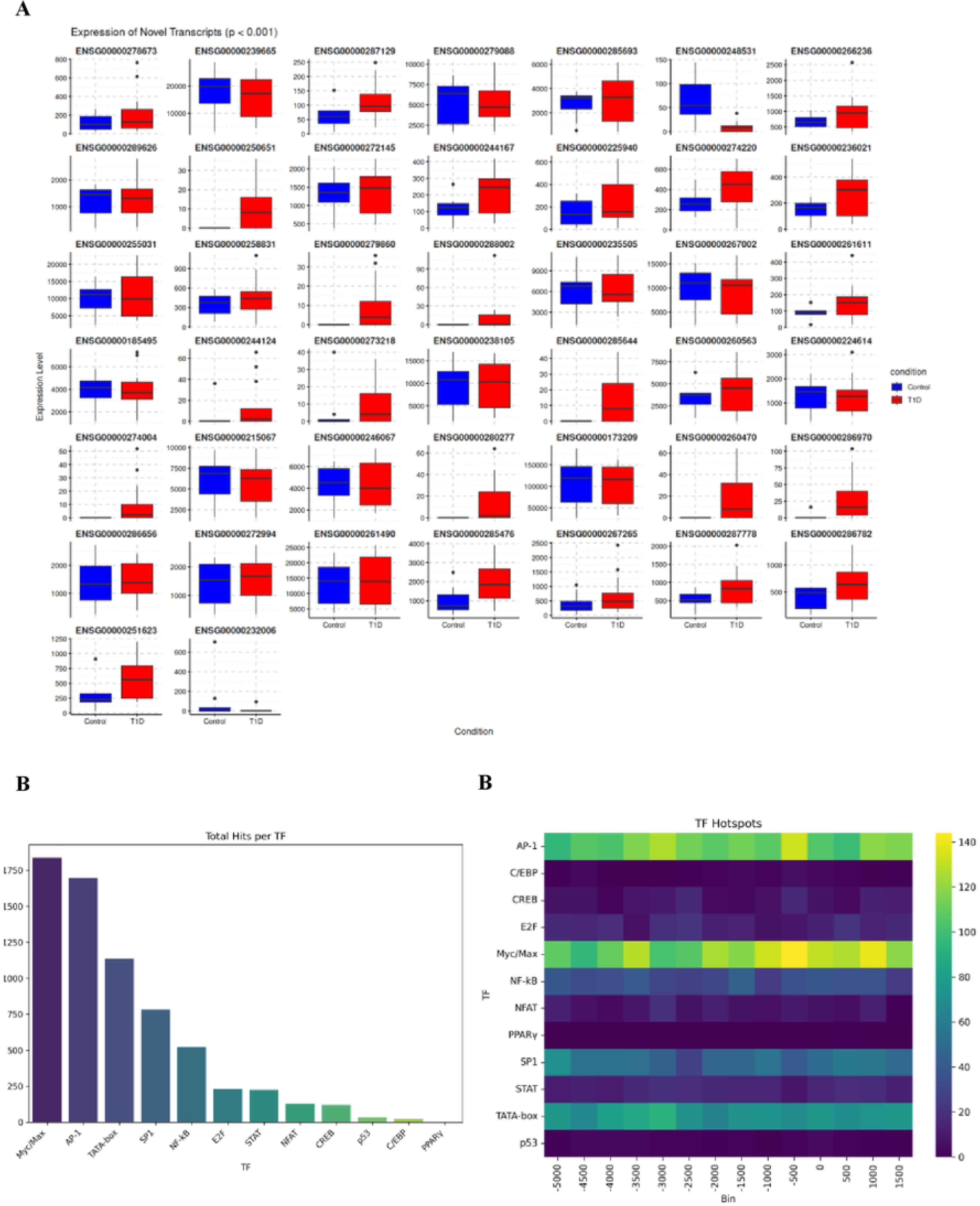
Transcription factor (TF) binding patterns in novel T1D transcripts. This figure characterizes TF regulatory dynamics in novel T1D transcripts. **Panel (A)** highlights 44 highly significant (p ≤0.001) uncharacterized novel transcripts lacking prior functional or disease associations, proposed as potential T1D biomarkers. **(B)** Illustrates the frequency of TF motif matches across genomic regions, derived from regex-based motif searches and visualized via a Seaborn barplot, emphasizing dominant motifs. (C) HeatMap of TF binding density across promoter-adjacent regions (-5 kb upstream to +2 kb downstream of transcription start sites) using a heatmap, revealing spatial regulatory hotspots. Together, these panels integrate positional, quantitative, and functional insights into TF-driven transcriptional dysregulation in T1D.

### Upstream motif profiling analysis reveals distinct TF binding patterns in T1D-associated transcripts including in the newly identified transcripts

To explore transcriptional regulation underlying differentially expressed genes in T1D, we performed upstream motif profiling across significantly up- and down-regulated gene sets. Using exact IUPAC-encoded TF motif matching within a ±5 kb region of each transcript, we observed a markedly higher frequency of motif occurrences in up-regulated genes compared to down-regulated genes (Fig 6A). Notably, the Myc/Max motif showed a nearly fivefold increase in up-regulated transcripts, followed by AP-1, SP-1, TATA-box, and NF-κB, each showing approximately fourfold enrichment (Fig 6A). These trends were consistently reflected in motif density heatmaps, which highlighted enriched TF binding hotspots in up-regulated genes relative to down-regulated counterparts (Fig 6B). To further characterize promoter activity in novel transcripts, we scanned the –5 kb upstream to +2 kb downstream regions surrounding their transcription start sites. Exact motif matches were quantified and visualized as bar plots (Fig 7A). Myc/Max showed the highest number of binding site hits, followed by AP-1, TATA-box, SP-1, and NF-κB, suggesting potential regulatory roles of these TFs in modulating novel gene activity. Heatmap analysis revealed distinct TF-specific hotspot distributions within the scanned regions (Fig 7B), indicating a complex and varied regulatory landscape at novel promoters.

## Discussion

Our blood-based transcriptomic survey of new-onset T1D patients versus matched controls uncovered 5,840 DEGs (1,916 stringent; p < 0.05, |log₂FC| > 2), and PCA of rlog counts confirmed a clear T1D signature (PC1 = 54 %, PC2 = 12 %). These findings echo extensive gene expression alterations reported in pancreatic tissues and islets, where chronic inflammation and innate immune activation drive β-cell damage (22) and early downregulation of immunity-related genes is observed within the first year post-diagnosis (24), reflecting dynamic molecular shifts in T1D. GWAS analyses by Pociot and Lernmark (42) further underscore genetic-environmental interactions driving immune and β-cell defects, consistent with our findings. The miRNA-mediated immune targeting of β-cells (24, 43) and Ruegsegger et al.’s work on intrinsic islet metabolic deficits (44) support a dual model where immune-mediated destruction coexists with inherent β-cell vulnerabilities. Collectively, these studies reinforce that T1D arises from a confluence of genetic predisposition, environmental triggers, immune hyperactivity, and metabolic impairment, culminating in progressive β-cell failure. Our data contextualize these mechanisms, emphasizing their systemic manifestation in early disease stages.

Pathway enrichment revealed both anticipated and novel perturbations. Immune processes such as lymphocyte differentiation, macrophage activation and cytokine signaling dominate (45, 46), while β-cell and insulin-secretion pathways highlight emerging metabolic stress. Unexpectedly, neurodegenerative disease pathways (general neurodegeneration, Alzheimer’s, ALS, Huntington’s) were enriched (47–50), suggesting shared oxidative-stress and mitochondrial dysfunction mechanisms. GO analysis also uncovered dysregulated GTPase regulator and serine/threonine kinase activities (51), and disruptions to cytoskeletal components and focal adhesions (52), linking T1D to broader cellular-architecture and signaling defects.

Our integrative TF-occupancy profiling placed Myc/Max, AP-1, SP-1, TATA-box and NF-κB motifs at the heart of T1D transcriptional control with pronounced motif enrichment in upregulated genes implicating immune-inflammatory pathways (Figure 6). Striking position clustering of these motifs near transcription start sites (TSS) of novel transcripts including highly up and down-regulated, and 44 unannotated suggests undiscovered regulatory elements (Supplement information_3, 4 and 5 respectfully). Myc/Max dominates with consistent genomic positioning, indicating direct control over novel transcripts. Spatial hotspot mapping (Figure 7) reveals promoter-centric, TF-specific regulatory coordination. These TF hotspots, particularly near novel transcripts, likely orchestrate coordinated gene-network shifts and represent high-value intervention points.

In addition, our TF-Gene interaction and signaling-network of these 211 genes revealed ∼ 72 significantly co-expressed (Supplment_Figure 1) interacting factors and gene diseases association using network-analyst analysis of these revealed 13 seed genes, identified as central or highly connected within the network and play significant role in the biological processes or diseases (Supplement_Figure 2). Understanding these networks and linking genes to diseases might help revealing how alterations might contribute to disease states and in identifying potential biomarkers and therapeutic targets (53, 54).

To our knowledge, this study is the first to deliver a systematic, quantitative map of transcription factor occupancy across Type 1 diabetes-specific transcripts, laying the groundwork for TF-targeted therapeutic strategies. Nevertheless, these in silico predictions demand experimental validation. Chromatin immunoprecipitation (ChIP) assays in relevant human cell lines will be critical to confirm direct TF-DNA interactions at the loci of interest. In parallel, studies in T1D animal models should determine whether the same transcriptomic changes and TF binding patterns occur in disease-relevant tissues. Because our data derive from whole-blood RNA, it will be important to pinpoint the cellular and tissue origins of these expression shiftswhether in pancreatic islets, pancreatic-draining lymph nodes, or other immune compartmentsto fully understand their role in pathogenesis. Finally, pharmacological modulation of the key TFs identified here, tested in murine T1D models, will reveal whether restoring or inhibiting their activity can prevent disease onset or progression, thereby advancing novel, mechanism-driven therapies for T1D. Interestingly, the insulin receptor signaling cascade emerged as a central axis in our data, alongside impaired regulatory T-cell function and altered cytokine signalingkey mediators of β-cell apoptosis (36–38, 39–41). Although these pathways are classically studied within pancreatic islets, similar TF-driven programs may operate in lymphoid tissues, where immune cells originate and become activated. Given that erythrocytes lack nuclei, our whole-blood transcriptome primarily reflects leukocyte biology. Dissecting which specific immune subsets drive these changes is therefore critical. The extensive expression shifts we observe likely stem from altered circulating immune cell proportions and activation states: expanded cytotoxic CD8⁺ T cells and pro-inflammatory CD4⁺ Th1/Th17 cells underpin many up-regulated inflammatory transcripts, while diminished FOXP3⁺ regulatory T-cell signatures indicate a loss of immune suppression. Concurrently, monocytes, dendritic cells, and natural killer cells contribute innate inflammatory signals, and elevated B-cell– associated transcripts underscore enhanced antigen-presentation activity. Collectively, these leukocyte-centered changes in cellular composition and activation state explain the peripheral transcriptomic remodeling characteristic of new-onset T1D.

Despite these insights, our study has several limitations: a modest sample size (n = 21), reliance on secondary whole-blood datasets that may not capture tissue-specific pathology or diverse patient backgrounds, and potential divergence between blood and pancreatic islet transcriptomes. To address these gaps, future work must expand and diversify patient cohorts, including paired blood and pancreatic samples to validate our DEGs and TF signatures across populations; employ targeted qPCR and chromatin immunoprecipitation assays to confirm TF–DNA interactions and novel transcript expression in relevant cell types; integrate complementary multi-omics layers (e.g., ATAC-seq, proteomics, metabolomics) to connect TF occupancy with downstream functional effects; and leverage single-cell and spatial transcriptomic approaches to resolve the specific leukocyte and islet cell subpopulations driving these transcriptomic shifts. Such comprehensive validation and refinement will strengthen our mechanistic models and accelerate the translation of TF-centric diagnostics and personalized therapies for Type 1 diabetes (55, 56).

In conclusion, our integrative platform delivers the first comprehensive, genome-wide quantification of transcription factor occupancy across Type 1 diabetes–specific transcripts, uncovering novel regulatory circuits that underpin disease onset. By illuminating uncharacterized transcripts alongside their key TF drivers, this work generates a rich set of candidate biomarkers and therapeutic targets. Ultimately, the ability to monitor TF dynamics with high precision in patient-derived samples holds transformative promise for personalized diagnostics and TF-focused interventions in T1D.

**Table 1.** Differential gene expression profiles and transcriptomic signatures T1D. This table summarizes the identification of 5,840 differentially expressed genes (DEGs) in T1D (4,800 upregulated; 1,040 downregulated). Applying stringent thresholds (*p* <0.05, |log₂FC| >2), 1,916 robust DEGs were retained, including 368 significantly upregulated and 96 significantly downregulated genes. Notably, 533 transcripts exhibited strong upregulation (log₂FC >5; 221 significant) and 44 showed marked downregulation (log₂FC <–5; 19 significant). The analysis further identified 211 novel transcripts (164 upregulated, 47 downregulated) with no prior disease associations, underscoring their potential as T1D-specific biomarkers.

| Up/down-regulated | DEGs categories | Criteria ($\log_2$ Fold chage, p-value) | Gene Number counts | Total (Novel transcriptomes, (p <0.05) | Total DEGs, (p <0.05) |
| --- | --- | --- | --- | --- | --- |
| **Up regulated** | **Up-regulated** | $\log_2FC > 2$ | 4800 | **211** | **1916** |
| | **Significant up-regulated** | $\log_2FC > 2, (p < 0.05)$ | 368 | | |
| | **Highly up-regulated** | $\log_2FC > 5$ | 533 | | |
| | **Significant highly up-regulated** | $\log_2FC > 5, (p < 0.05)$ | 221 | | |
| | **Significant novel transcript** | $(p < 0.05)$ | 164 | | |
| **Down regulated** | **Down-regulated** | $\log_2FC < -2$ | 1040 | | |
| | **Significant down-regulated** | $\log_2FC < -2, (p < 0.05)$ | 96 | | |
| | **Highly down-regulated** | $\log_2FC < -5$ | 44 | | |
| | **Significant highly down-regulated** | $\log_2FC < -2, (p < 0.05)$ | 19 | | |
| | **Significant novel transcript** | $(p < 0.05)$ | 47 | | |

## Declaration statement

### Availability of data and materials

All data generated or analyzed during this study are included in this published article. Related processed intermediary data files and automated python and R scripts could be provided upon reasonable request.

### Competing interests

The authors declare that there are no conflicts of interest.

## Acknowledgments

The authors thank Dr. Bobbie-Jo M Webb-Robertson and colleagues for generously providing access to their raw sequencing data, which were critical to this study. Their original work, “RNA Splicing Events in Circulation Distinguish Individuals With and Without New-onset Type 1 Diabetes” (The Journal of Clinical Endocrinology & Metabolism. 2024 Sep 10: dgae622, 2025), enabled the comparative analyses presented here. We acknowledge the importance of open data sharing in advancing scientific discovery and commend their contribution to the field of type 1 diabetes research.

## Notes

### Competing Interest Statement

The authors have declared no competing interest.

